# Assessing preferences for adult vs juvenile features in young animals: newly-hatched chicks spontaneously approach red and large stimuli

**DOI:** 10.1101/2023.02.17.528933

**Authors:** Laura Freeland, Vera Vasas, Josephine Gomes, Elisabetta Versace

## Abstract

Young precocial birds benefit from staying close to both their mother and siblings, while prioritising adults, which provide better care. Which features of the stimuli are used by young birds to prioritise attachment to adults over siblings is unknown. We started to address this question in newly hatched domestic chicks (*Gallus gallus*), focusing on their spontaneous preferences for visual stimuli that systematically vary between adult and juvenile chickens: size (larger in adults than in juveniles) and colour (darker and redder in adults than in juveniles). Overall, chicks at their first visual experience, that had never seen a conspecific beforehand, were most attracted to the red and large stimuli (two adult features) and interacted with red stimuli more than with yellow stimuli. When tested with red large vs. small objects (Exp. 1), chicks preferred the large shape. When tested with yellow large and small objects (Exp. 2), chicks did not show a preference. These results suggest that the combination of size and colour form the predisposition that helps chicks to spontaneously discriminate between adult and juvenile features from the first stages of life, in the absence of previous experience.

## 1. Introduction

To benefit from protection and acquire relevant information, young social animals should preferentially direct their attention towards adult conspecifics. Indeed, in many species we observe brood movements led by the mother, with juveniles following the adult (Fig. 1). To help group cohesion, chickens and other precocial birds have evolved a fast-learning mechanism called imprinting [1–3], that enables them to quickly learn the specific features of social partners and stay close to them. After a brief imprinting exposure with the mother hen (in experimental settings also with another natural or artificial moving object such as the ethologist Konrad Lorenz [4], a plastic cylinder [5–7] or even a shape displayed on a computer monitor [8–10]), chicks develop attachment for the imprinting object, follow it when moves and produce distress calls when they are separated from it. However, while young birds are exposed to the hen, they are often exposed to the siblings of the same batch too. In wild fowls, batches consist of 4-6 eggs [11,12], that hatch at around the same time. This produces an interference between filial imprinting directed toward the mother hen, and other salient conspecifics [13]. Interestingly, young chicks do imprint on their siblings, an adaptive behaviour that supports group cohesion and predator avoidance [14–18]. Despite these benefits, it would be maladaptive for chicks to preferentially imprint on their siblings, because the mother hen can provide more warmth, protection from predators and relevant information compared to the inexperienced siblings. What mechanisms enable young precocial birds to preferentially orient to and then imprint on adult hens vs. young siblings is unknown.

**Figure 1.**
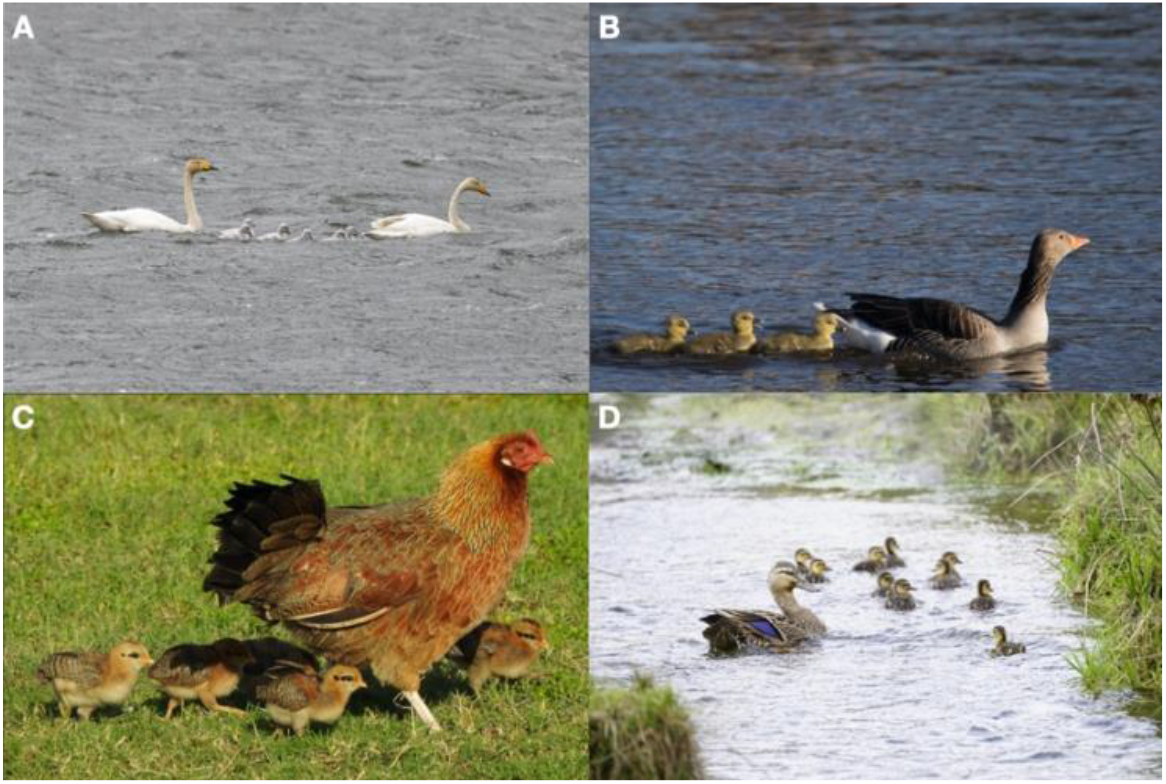
In precocial birds, juveniles imprint on and follow their mother soon after hatching. (A) Whooper swan (*Cygnus cygnus*) (B) Graylag goose (*Anser anser*) (C) Domestic chicken (*Gallus gallus*) (D) Mallard (*Anas platyrhynchos*). Image credits: Macaulay Library at the Cornell Lab of Ornithology (Mason Flint, [ML505596021](https://macaulaylibrary.org/asset/505596021); Roland Pfeiffer, [ML442529161](https://macaulaylibrary.org/asset/442529161); Joel Gilb [ML198044131](https://macaulaylibrary.org/asset/198044131); Ulises Cabrera Miranda, [ML494249871](https://macaulaylibrary.org/asset/494249871)).

We hypothesise that spontaneous preferences available in the absence of previous experience – *predispositions* [19–22]) – might help chicks to preferentially orient towards adult vs. juvenile animals, and therefore to preferentially imprint on adults. We start addressing this question in newly hatched domestic chicks (*Gallus gallus*), focusing on their spontaneous visual preferences for cues associated with adults in terms of size and colour. We investigated visual stimuli because extensive evidence shows that newly hatched chicks primarily rely on vision [1,8]. At the beginning of life, chicks use low-level visual features to orient their initial approach preferences [reviewed in 19,20,22]. These predispositions include spontaneous preferences for the neck and head region [23–26], biological motion [27], upward movement [28], changes in speed [29,30], combination of colour and biological motion [31], size [32,33] and colour [e.g. 31, reviewed in 34]. It is not known whether predispositions are preferentially directed to adult-like features.

Regarding size, we reasoned that adult chickens are consistently larger than juveniles (Fig. 1), as in other vertebrate species. Accordingly, we hypothesized that the larger size may be a cue used by young animals to preferentially attach to their parents. In line with this idea, newly hatched chicks have been shown to react more to multicoloured balls with a diameter of 10-20 cm, being less responsive to diameters of 5 or 25+ cm [32]. This pattern is suggestive of stronger preference for objects with a size similar to that of the mother hen vs. objects with a size similar to that of newly hatched chicks. However, this work did not test the relative preference for the same proximal stimulation (e.g., same area) distributed between larger vs. smaller individuals. More recently, chicks previously imprinted with bidimensional objects have been tested for their preferences for objects of different size, colour and shape after imprinting [33]: chicks preferred larger numerosities of similarly sized objects irrespective of the number of stimuli they had been imprinted on, choosing larger volumes rather than familiar numerosity when continuous features were controlled for. This work left open the question on the spontaneous preferences of chicks for objects that differ in colour and size, before they have been imprinted. Our experiments address this knowledge gap, focusing on the spontaneous approach responses for a single large vs five small objects (controlled for colour area), before imprinting takes place. We predicted that chicks should have a spontaneous preference for the single large object.

We also analysed colour preferences, another feature that can be used by chicks to discriminate between adults and juveniles. Previous work has shown that colour is central for behavioural choices in young chicks: for instance, colour is more important than shape in establishing imprinting preferences [9,35,36]. Chicks are able to process fine differences in colours from the first days of life, as shown in pecking/foraging tasks [37–39]. Interestingly, previous experiments [40] showed that red colour paired with biological motion is more effective than yellow colour matched with biological motion in eliciting chicks’ preferences. To clarify whether colour preferences reflect systematic differences between hens and juveniles in the wild, we investigated the spectral reflectance of the red junglefowl (*Gallus gallus*), the closest living relative of the ancestral wild chicken before domestication [41]. We investigated systematic colour differences in feather colour and head colour between adults and juveniles using spectral reflectance measurements of stuffed bird skins from the Natural History Museum collections and pictures from the Macaulay library (see Fig. 2). Previous studies have shown that the head is a crucial body region for driving preferential approach choices [23,25,42,43], hence our focus on this area. As detailed below, we found that both hens and chicks display yellowish and brownish colours, but hens are darker (Fig. 2A). Moreover, in hens the head region is more reddish than in chicks and juveniles (Fig. 2D). Based on these observations, we predicted that young chicks should spontaneously prefer darker and more reddish colours, as a strategy to preferentially approach adult hens.

**Figure 2.**
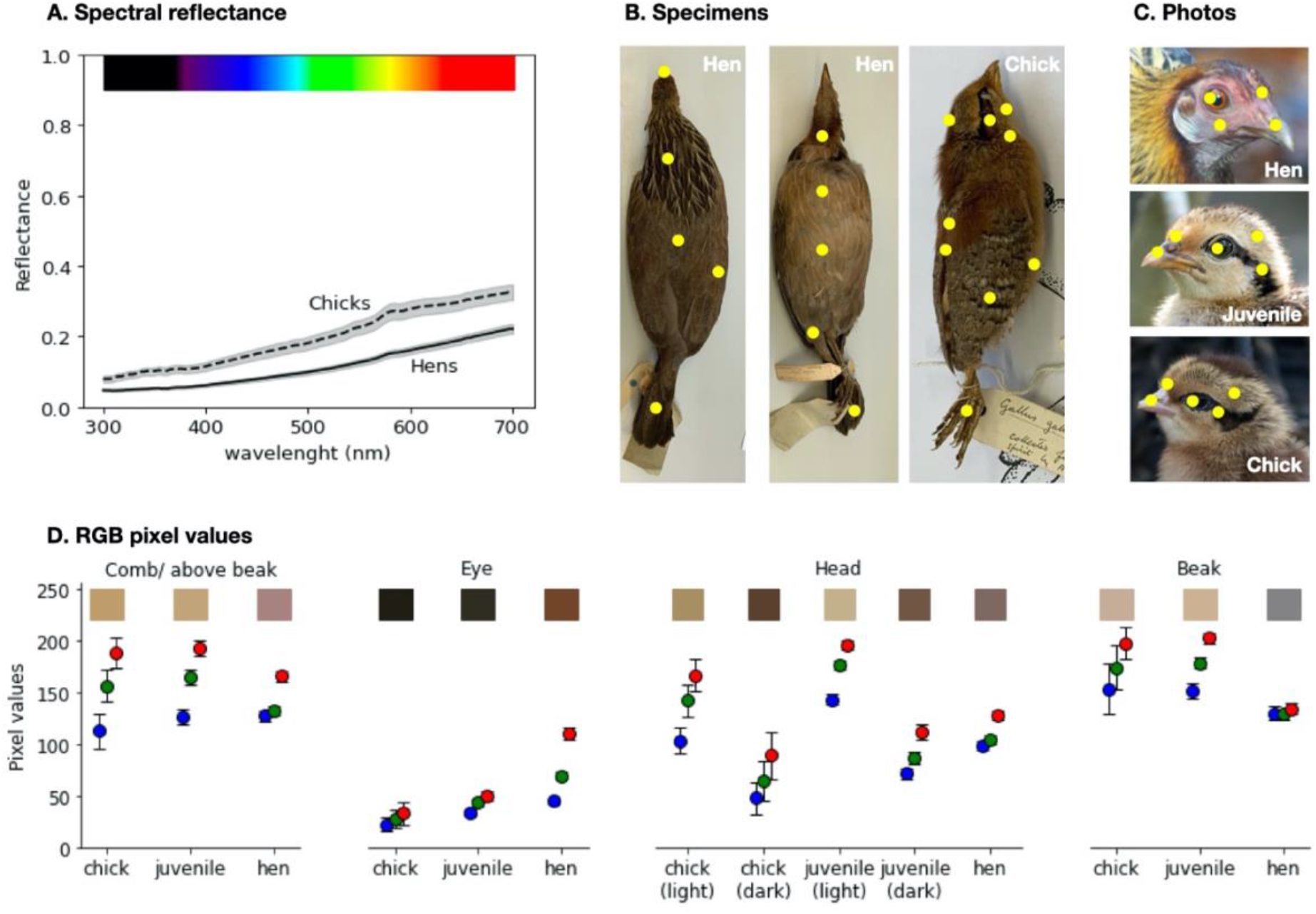
Colours of red junglefowl (*Gallus gallus spadiceus*) hen and chicks. (A) The average spectral reflectance of the feathers of hens and chicks. The shadow indicates the standard error at each wavelength. (B) Example specimens of a hen and a chick of the red junglefowl. Yellow dots mark the body parts sampled with spectrophotometry. From the hens, we sampled the chest, neck underside, belly, upper leg, feet, tail underside, back, tail, neck from above, wing and beak areas. From the chicks, we took samples from the centre, side and underside of the head, from the underside of the neck, belly, feet, wing and from the dark and light stripes on the back. (C) Examples photos of the red junglefowls. Yellow dots mark the body parts sampled from photos: the comb (when present) or the area above the beak, the eye, the head (two measurements for the bicoloured chicks and juveniles) and the beak. Image credits: Macaulay Library at the Cornell Lab of Ornithology (James Hully, [ML471555411](https://macaulaylibrary.org/asset/471555411); Kiri Zhang, [ML478986371](https://macaulaylibrary.org/asset/478986371); Andreas Boe, [ML200531451](https://macaulaylibrary.org/asset/200531451)). (D) Mean ± SEM of RGB values of the pixels taken from photos of red junglefowl hens, juveniles and chicks. Red, green and blue dots indicate the red, green and blue components of the RGB values; the coloured rectangles illustrate the average colour.

To test our predictions, we hatched chicks in darkness, and then presented them with stimuli of different size and colour using computer animations [see also 8,9]. At their first visual experience, chicks were given a two-choice test between one large shape vs. five small shapes with same total area. In Experiment 1, shapes were red, hence the large stimulus had two “adult-like” features, i.e. size and colour. In Experiment 2, shapes were bright yellow, hence the large stimulus has only one “adult” feature, i.e. size. If chicks use size and colour to preferentially orient towards adults, we expected a stronger preference for the larger stimuli in both experiments, with a stronger effect in Experiment 1 compared to Experiment 2, due to the presence of two adults features in Experiment 1.

## 2. Methods

### 2.1 Colour assessment of feathers and head in hens and chicks

Spectral reflectance measurements were taken at the Bird Collection of the Natural History Museum (Tring, UK) using an AvaSpec-2048 spectrophotometer (Avantes) with an AvaLight-DHS (Avantes) light source on stuffed bird specimens. We measured eleven different areas on four specimens of hens, and nine differently areas on three specimens of young chicks of red junglefowl (*Gallus gallus spadiceus*), see Fig. 2B. Comparing hens and chicks, we observed that the average reflectance curves have the same shape (Fig. 2A), with a gradual increase in reflectance with larger wavelengths (characteristic of pheomelanin), resulting in the yellowish-brown appearance of the specimens (Fig. 2B-C). The skin of chicks consistently reflects more light at every wavelength, appearing brighter than hens’ skin.

In stuffed specimens the soft parts of the body, including the head region, dry out; for this reason, we could not assess their original colour. To compare the colour of the head of chicks and hens, we used photos of wild red junglefowl taken from the Macaulay library. We extracted the RGB pixel values of 60 hens, 43 juveniles and 9 newly-hatched chicks from comparable locations on the head (see Fig. 2C). None of the colours were saturated, but the hens appear slightly reddish in the comb/beak, eye and head areas (with higher red than blue/green RGB values). The chicks and the juveniles have two main colours in these areas: the comb/beak and the lighter feathers on the head are slightly yellowish (with higher red/green than blue RGB values) and brighter than in hens; the eye is black (without the orange band present in hens) and there is a dark brown stripe on the head [44]. The beak is light brown in the chicks/juveniles and a darker grey in hens. Overall, the head regions of hens is darker and more reddish than those of chicks.

### 2.2 Predisposition experiments Subjects and rearing conditions

After standard incubation in darkness (37.7 °C and 40-60% humidity), chicks hatched in individual compartments and were carried to the experimental room for testing in an opaque box. For incubation and hatching we used a FIEM incubator S140ADS and hatchery H316DS. We tested domestic chicks (*Gallus gallus*) of the Ross 309 strain: 51 chicks (25 females, 26 males) in Experiment 1 (red stimuli), and 62 chicks (27 females, 35 males) in Experiment 2 (yellow stimuli), see Table 1 for details. Chicks were tested within 14 hours after hatching [30]. Experiments adhere to National regulations on animal research, and have been approved by the University ethics committee (AWERB) and Home Office (PP5180959).

**Table 1.** Number of chicks used in the experiments.

|  | Total collected | Low quality videos | Total chicks analysed | No choice |
| --- | --- | --- | --- | --- |
| Exp 1. | 59 (28♀, 31♂) | 8 (3♀, 5♂) | 51 (25♀, 26♂) | 4 (1♀, 3♂) |
| Exp 2. | 69 (28♀, 41♂) | 7 (1♀, 6♂) | 62 (27♀, 35♂) | 22 (5♀, 17♂) |

#### Apparatus and stimuli

The experimental apparatus (Fig. 3) consisted of a wooden box (90×60×52 cm) with non-slip black floor, white walls and two monitors at each end displaying at 120 fps. The apparatus was divided into three virtual regions: the centre (54×60 cm) and two side choice areas (18×60 cm). A videocamera located on top of the arena recorded the entire session. The experiments were recorded at 10 frames per second, at the resolution 1280×720 pixels.

**Figure 3.**
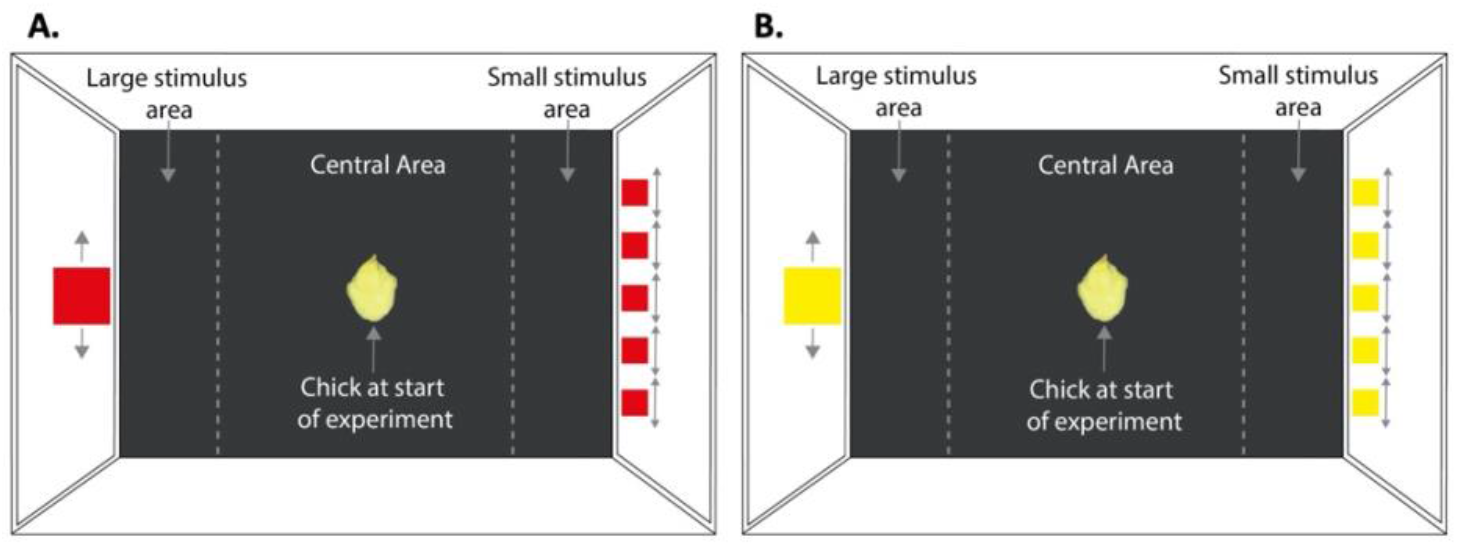
Experimental setup in the Experiment 1 (A) and Experiment 2 (B). The arena was virtually divided into a central “no-choice” area where the chick was located at the beginning of the experiment, and two “side choice” areas. The centroid position of the chick was recorded and automatically tracked. In Experiment 1 (A), the chicks were given a choice between a large red square vs. five small red squares. To enhance approach responses, the stimuli moved horizontally, right and left, for an equal duration and distance on each side, as indicated by the arrows. In Experiment 2 (B), the chicks were given a choice between a large yellow square vs. five small yellow squares.

The stimuli were displayed on two 24” Asus VG248QE monitors. A large shape (111.8 mm square and total area of 125 cm^2^) was shown on one monitor, and five small shapes (50 mm squares) on the opposite. The surface area of the single large shape was equal to the five small shapes. In Experiment 1, the stimuli were red (RGB: 255, 0, 0; Fig. 3A), in Experiment 2, yellow (RGB: 255, 255, 0; Fig. 3B). To enhance approach responses, the stimuli moved horizontally. The larger stimulus moved 50 mm back and forth for five seconds while each smaller stimulus moved, one after the other, by 50 mm, so the objects moved for an equal duration and distance on each side. The stimuli remained still for a further two seconds, then a white screen was shown for two seconds. Following the white screen, the stimuli reappeared in a different position. Five different locations were played resulting in a total play through of 60 seconds. The right-left position of the stimuli in the arena was counterbalanced between subjects.

#### Procedure

At the start of the experiment, chicks were gently placed in the centre of the apparatus with the beak facing the long wall. In this way, through their lateral vision, subjects could see both stimuli at the same time, before deciding which stimuli to approach. Chicks were recorded for 20 minutes by a camera mounted above the centre of the arena. Chicks were free to move in the apparatus.

#### Data analysis

The chicks’ position in the arena was tracked automatically using DeepLabCut [45]. For a recording to be included in the analysis, 90% of the video frames must have had an accuracy likelihood of 0.9 or greater [see 28]. In Experiment 2, in 6 videos the recording was not clear enough to identify the stimuli, and we discarded those data. See table 1 for details.

To analyse the chicks’ initial preferences, we have recorded the first choices, defined as the centroid of the chick entering either stimulus area. Significant differences in the number of first choices for large vs. small stimuli were assessed using binomial tests. The difference in latency to first approach (i.e., time elapsed until the chick made its first choice) was assessed using Mann-Whitney U tests, as latency did not match assumptions for parametric tests.

To analyse the chicks’ preference for the large stimulus, we calculated a preference index as

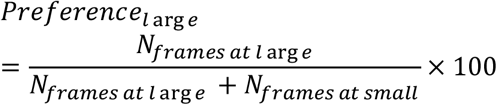

where 1 indicates a complete preference for the large stimulus, 0 a complete preference for the small stimuli and 0.5 indicates no preference, in line with previous research [9,46].

Time was divided into 20 one-minute time bins in which the preference of each chick was calculated for each minute. As the preference index did not match assumptions for parametric tests, we used Friedman tests to determine if the chicks’ preference changed over time. The bins were then averaged over across each experiment to calculate the mean preference for the large stimulus. We used one-sample Wilcoxon tests to establish if the preference for the large object deviated from chance level (0.5) and if the preference was different in the red vs. yellow experiment.

To compare the rate of subjects that approached the stimuli between Experiment 1 and 2 we used a Chi-square test. To compare the responsiveness to the stimuli across experiments we analysed the total time spent in the centre using Mann-Whitney U tests. For all tests we set alpha to 0.05.

## 3. Results

In Experiment 1, 47 out of 51 tested chicks (91%) approached at least one stimulus. Chicks preferred to approach first the larger stimulus (35 vs. 11 chicks, 76%; p<0.001). The preference for the large stimulus was different across time bins (χ^2^=56, p<0.001). The difference was driven by the time it took the chicks to move towards their chosen stimulus and it disappeared after the first 5 minutes (bins 5-20: χ^2^=3.49, p=0.479). Out of the 20 one-minute time bins, 17 showed significantly higher preference for the large stimulus (all except bins 1, 13 and 14, Fig. 3). As the preference remained at or above chance level (50%) throughout the test (Fig. 4), we collapsed the data across time bins. The mean preference for the large stimulus across the whole experiment was significantly higher than random choice (Fig. 5, M± SEM=0.666±0.045, median=0.805; V=243.5, p=0.001). Overall, chicks preferred the single red large stimulus, as expected.

**Figure 4.**
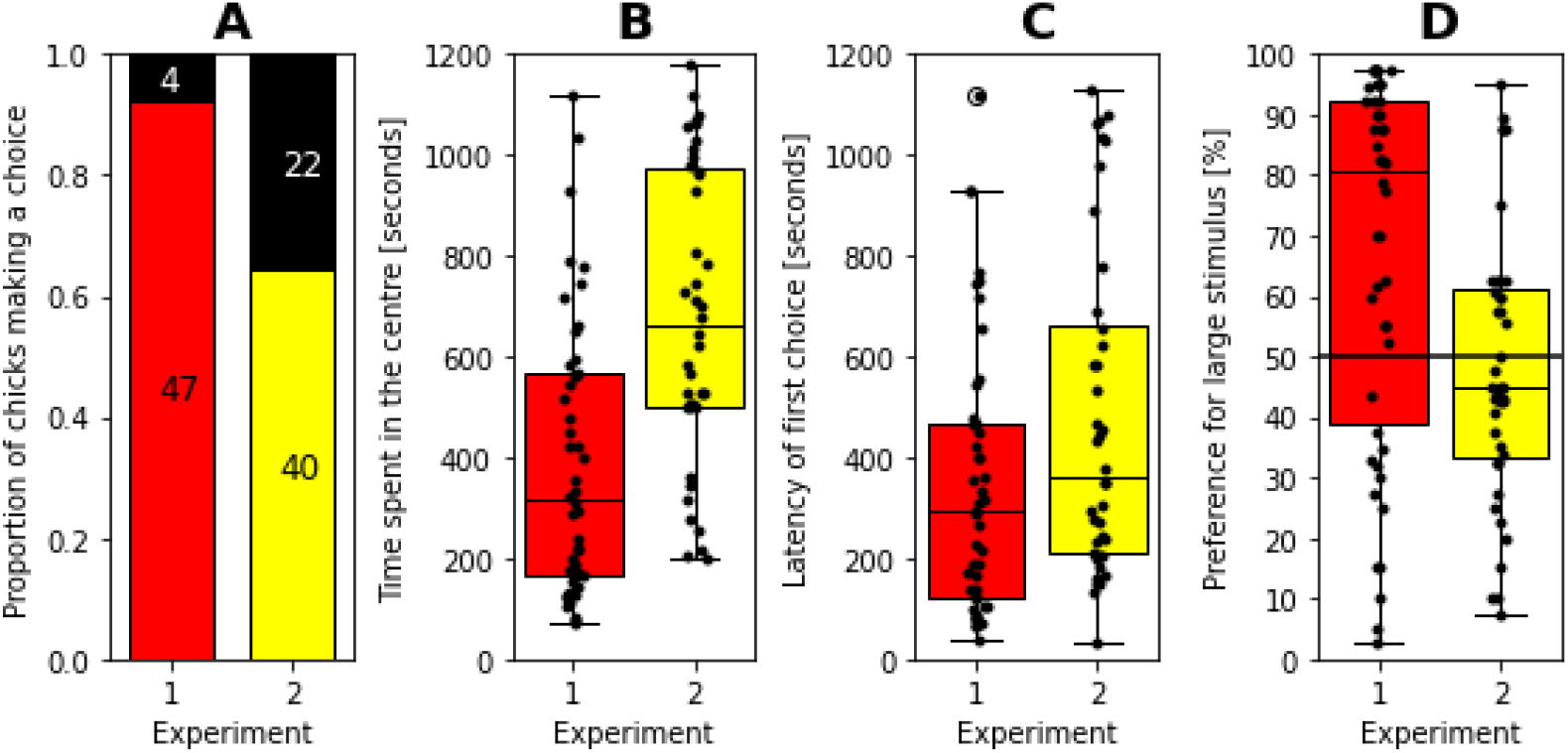
Comparisons of Experiment 1 (red stimulus) and Experiment 2 (yellow stimulus). The subplots show the (A) proportion of the chicks that made any choice in the 20 minutes of the experiment and the number of chicks in each category; the (B) the time the chicks have spent in the centre area; (C) the time elapsed until they made their first approach to any of the stimulus areas and (D) their preference for the large stimulus. In the subplots B-D, dots indicate the individual data points, and the boxplots show the median, first quartile to the third quartile of the data, the most extreme non-outlier points and the outliers.

**Figure 5.**
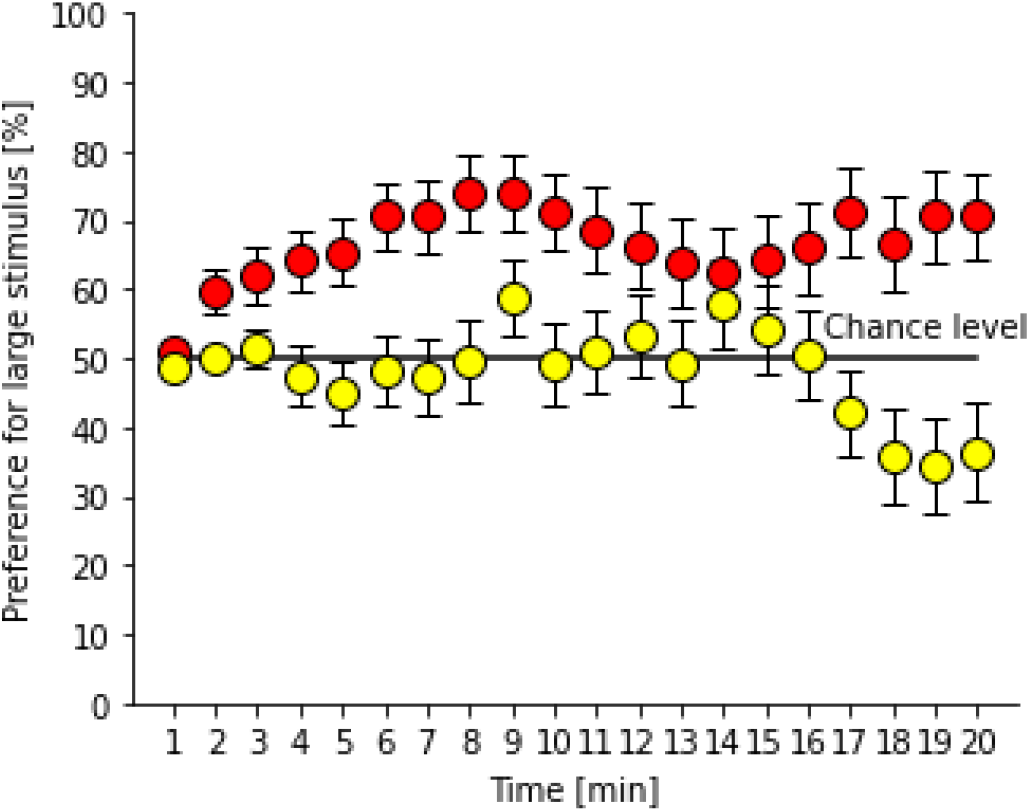
Preference for the large stimulus in Experiment 1 (red) and Experiment 2 (yellow), over time. Dots indicate the average preference of the chicks for that minute and error bars show the standard error of the mean.

In Experiment 2, out of 62 tested chicks, 22 remained in the centre, and only 40 (65%) approached either of the experimental stimuli. Out of these, more chicks first approached the smaller stimuli than the large stimulus; however, the difference was not significant (24 vs. 16, 60%; p=0.268). The preference was not significantly different across time (χ^2^=8.6, p=0.474). Looking at time bins individually (Fig. 4), two bins (18 and 19) reached the significance threshold (p = 0.035 and 0.024); however, neither was significant after correction for multiple comparisons (using Bonferroni correction, the overall alpha level of 0.05 would require a p < 0.0025 for an individual hypothesis). As the preference remained between chance and preference for the small objects throughout the test (Fig. 5), we collapsed data across time bins. The mean preference for the large stimulus trended for a preference for small objects (Fig. 5, M ± SEM=0.480 ± 0.035, median=0.450; V=363.0, p=0.527). Overall, we have not found evidence for size preference when the stimuli were shown in yellow.

Comparing the two experiments, we have found differences in the number of chicks that made a choice. Chicks in Experiment 1 (red stimuli) were more likely to make a choice than in Experiment 2 (yellow stimuli) (χ^2^=10.56, p=0.001). Chicks that approached the stimuli were significantly slower in Experiment 2 (yellow stimuli), (Fig. 5, latency to first approach in Exp. 1, M ± SEM: 345 s ± 39, latency in Exp.2, M ± SEM=481 s ± 51; U=677.0, p=0.036) and accordingly spent more time in total in the centre area (Fig. 5, red stimuli M ± SEM=389±40 seconds, yellow stimuli M± SEM=682±46 seconds; U=420, p<0.001). Taken together, these findings indicate a stronger responsiveness to red than yellow stimuli.

## 4. Discussion

Which features of the stimuli are used by young animals to prioritise attachment to adults over juveniles has not been investigated, despite the advantage for younger animals to receive parental care from adult animals. We addressed this question by testing the spontaneous preferences of newly hatched domestic chicks (*Gallus gallus*). We investigated the attractiveness of size (large vs. small) and colour (red vs. yellow), two easily discernible features that differ between adults and juveniles. We hypothesised that by focusing on larger size and colours associated with adults, rather than on smaller size and colours associated with juveniles, chicks could reliably prioritise affiliative responses towards adults.

In vertebrate species, adults grow larger than young animals, hence a larger size is a univocal feature associated with adulthood. Colour differs between species and developmental stages. To detect consistent colour differences between hens and chicks, we focused on the head region, that is important in driving chicks’ preferences [24,25,40,42]. We analysed stuffed specimens and pictures of the Red jungle fowl (*Gallus gallus spadiceus*). The Red jungle fowl is the closest relative to the ancestral wild chickens before domestication [41]. Overall, hens and chicks have similar spectral reflectance, but hens are darker and more reddish, compared the lighter and more yellowish chicks. Based on this, we tested chicks for their spontaneous preference to approach red or yellow displays or large of small size, controlling for the area and movement of visual stimuli presented.

When presented with large vs. small red visual stimuli (Exp. 1), visually inexperienced chicks had a spontaneous preference for the larger stimuli. This result is in line with previous findings for the most attractive size [47] and for preference for larger objects [33]. This spontaneous preference for the larger stimulus overcame the chicks’ previously observed preference for larger numerosities [33]. This result supports the hypothesis that size can draw a young chick’s attention towards adults before imprinting takes place.

When presented with large vs. small yellow visual stimuli (Exp. 2), the chicks had no significant preference. A preference for the yellow and small stimuli observed towards the end of the experiment was not significant after correction for multiple comparisons. Should a preference for lighter and smaller objects be confirmed, it would suggest that chicks have spontaneous expectations for the association of adult (large and red) vs. juvenile (small and yellow) features. Given the mounting evidence on crossmodal associations in chicks [46,48], this possibility should be further investigated.

Comparing the two experiments, the yellow stimuli were consistently less attractive. When yellow stimuli were presented, chicks spent significantly less time in the areas close to the stimuli, and significantly fewer chicks approached the stimuli. A lower interest for the yellow stimuli has been previously observed in experiments that didn’t manipulate the size of visual stimuli [49–51]. Further experiments should clarify to what extent this effect is driven by hue, brightness, a combination of both these colour features and the contrast with the background. Previous evidence on colour preferences in chicks in non-univocal, likely due to the fact that chicks have been tested with different methods, tasks, backgrounds, and at different ages [37,e.g. 49,50,52]. However, our results are in line with the consensus that red and to a lesser extent blue are consistently attractive colours for chicks [49–51], while green and yellow elicit weaker approach responses [37,50]. Interestingly, it has been found that chicks do imprint on yellow objects, but the process takes longer [53]. The longer exposure required to imprint on yellow objects may enable chicks to imprint on both adult and siblings, thus keeping group cohesion, and at the same time prioritise the affiliative responses towards the adults.

This first work on predispositions for features that indicate the presence of adult features adds to the growing body of evidence on spontaneous predispositions. Such predispositions support the adaptive responses of animal at the start of life predator avoidance [54], preferential attention to animate objects [19,22,55–57] and the imprinting on care-giving adults, as discussed here.

## Authors’ contributions

L.F: conceptualization, methodology, investigation, formal analysis, visualization, writing: original draft and writing: review and editing; V.V: methodology, software, formal analysis, investigation, data curation, validation, visualization, writing: original draft and writing: review and editing; J.G.: validation, formal analysis, writing review and editing; E.V.: conceptualization, project administration, supervision, writing: original draft and writing: review and editing. All authors gave final approval for publication and are accountable for the work performed therein.

## Conflict of interest declaration

We declare we have no competing interests.

## Funding

This work was supported by the Leverhulme grant RPG-2020-287 (E.V.) and the Royal Society Leverhulme Trust fellowship SRF\R1\21000155 (E.V.)

## Acknowledgements

We thank Shuge Wang for helpful discussions on this project, including video analysis. We are grateful for The Macaulay Library at the Cornell Lab of Ornithology for providing images.

## Notes

### Competing Interest Statement

The authors have declared no competing interest.

### Summary of Updates

credits images

## References

1. Bolhuis JJ. 1999 Early learning and the development of filial preferences in the chick. Behavioural brain research 98, 245–52.

2. McCabe BJ. 2019 Visual Imprinting in Birds: Behavior, Models, and Neural Mechanisms. Frontiers in Physiology 10. (doi:10.3389/fphys.2019.00658)

3. Vallortigara G, Versace E. 2018 Filial Imprinting. In Encyclopedia of Animal Behavior, pp. 1943–1948.

4. Lorenz K. 1937 The Companion in the Bird’s World. The Auk 54, 245–273.

5. Versace E, Regolin L, Vallortigara G. 2006 Emergence of Grammar as Revealed by Visual Imprinting in Newly-hatched Chicks. In: The Evolution of Language. In Proceedings of the 6th International Conference, Rome, 12-15 April, pp. 457–458.

6. Vallortigara G, Andrew RJ. 1994 Differential involvement of right and left hemisphere in individual recognition in the domestic chick. Behavioural Processes 33, 41–57. (doi:10.1016/0376-6357(94)90059-0)

7. Szabó E, Chiandetti C, Téglás E, Versace E, Csibra G, Kovács ÁM, Vallortigara G. 2021 Young domestic chicks spontaneously represent the absence of objects. bioRxiv, 2021.01.20.427266.

8. Versace E, Spierings MJ, Caffini M, ten Cate C, Vallortigara G. 2017 Spontaneous generalization of abstract multimodal patterns in young domestic chicks. Animal Cognition 20, 521–529. (doi:DOI 10.1007/s10071-017-1079-5)

9. Lemaire BS, Rucco D, Josserand M, Vallortigara G, Versace E. 2021 Stability and individual variability of social attachment in imprinting. Scientific Reports 11, 1–12. (doi:10.1038/s41598-021-86989-3)

10. Wood JN. 2015 Characterizing the information content of a newly hatched chick’s first visual object representation. Developmental Science 18, 194–205. (doi:10.1111/desc.12198)

11. Nicol CJ. 2015 The Behavioural Biology of Chickens. Oxfordshire OX10 8DE (UK): CABI.

12. Ali S, Anwar M, Rais M, Mahmood T. 2016 Breeding ecology of red jungle fowl (*Gallus gallus*) in Deva Vatala National Park, Azad Jammu and Kashmir, Pakistan. journal of Applied Agriculture and Biotechnology 1, 59–65.

13. Rivera M, Louder MIM, Kleindorfer S, Liu W, Hauber ME. 2018 Avian prenatal auditory stimulation: progress and perspectives. Behav Ecol Sociobiol 72, 112. (doi:10.1007/s00265-018-2528-0)

14. Lickliter R, Gottlieb G. 1988 Social specificity: Interaction with own species is necessary to foster species-specific maternal preference in ducklings. Developmental Psychobiology 21, 311–321. (doi:10.1002/dev.420210403)

15. Darczewska M. 1999 Peer Attraction in White Peking Ducklings (*Anas platyrhynchos*).

16. Vallortigara G, Andrew RJ. 1991 Lateralization of response by chicks to change in a model partner. Animal Behaviour 41, 187–194.

17. Bateson P. 1982 Preferences for cousins in Japanese quail. Nature 295, 236–237. (doi:10.1038/295236a0)

18. Zajonc RB, Wilson WR, Rajecki DW. 1975 Affiliation and social discrimination produced by brief exposure in day-old domestic chicks. Animal Behaviour 23, 131–138. (doi:10.1016/0003-3472(75)90059-7)

19. Rosa-Salva O, Mayer U, Versace E, Hebert M, Lemaire BS, Vallortigara G. 2021 Sensitive periods for social development: Interactions between predisposed and learned mechanisms. Cognition 213, 104552. (doi:10.1016/j.cognition.2020.104552)

20. Versace E, Vallortigara G. 2015 Origins of knowledge: Insights from precocial species. Frontiers in Behavioral Neuroscience 9, 338. (doi:10.3389/fnbeh.2015.00338)

21. Versace E, Ragusa M, Vallortigara G. 2019 A transient time window for early predispositions in newborn chicks. Scientific Reports 9, 18767. (doi:https://doi.org/10.1038/s41598-019-55255-y)

22. Versace, Martinho-Truswell. 2018 Priors in Animal and Artificial Intelligence: Where Does Learning Begin? Trends in cognitive sciences 1823.

23. Rosa-Salva O, Regolin L, Vallortigara G. 2010 Faces are special for newly hatched chicks: evidence for inborn domain-specific mechanisms underlying spontaneous preferences for face-like stimuli. Developmental Science 13, 565–77. (doi:10.1111/j.1467-7687.2009.00914.x)

24. Versace E, Fracasso I, Baldan G, Dalle Zotte A, Vallortigara G. 2017 Newborn chicks show inherited variability in early social predispositions for hen-like stimuli. Scientific Reports 7, 40296. (doi:DOI: 10.1038/srep40296)

25. Johnson MH, Horn G. 1988 Development of filial preferences in dark-reared chicks. Animal Behaviour 36, 675–683.

26. Rosa-Salva O, Mayer U, Vallortigara G. 2019 Unlearned visual preferences for the head region in domestic chicks. PLoS ONE 14, 1–15. (doi:10.1371/journal.pone.0222079)

27. Vallortigara G, Regolin L, Marconato F. 2005 Visually Inexperienced Chicks Exhibit Spontaneous Preference for Biological Motion Patterns. PLoS Biol 3, e208. (doi:10.1371/journal.pbio.0030208)

28. Bliss L, Vasas V, Freeland L, Roach R, Ferrè ER, Versace E. 2022 Newborn chicks prefer stimuli that move against gravity. (doi:10.1101/2022.07.13.499929)

29. Rosa-Salva O, Grassi M, Lorenzi E, Regolin L, Vallortigara G. 2016 Spontaneous preference for visual cues of animacy in naïve domestic chicks: the case of speed changes. Cognition 157, 49–60. (doi:10.1016/j.cognition.2016.08.014)

30. Versace E, Ragusa M, Vallortigara G. 2019 A transient time window for early predispositions in newborn chicks. Scientific Reports 9, 18767. (doi:10.1038/s41598-019-55255-y)

31. Miura M, Matsushima T. 2016 Biological motion facilitates imprinting. Animal Behaviour 116, 171–180. (doi:10.1016/j.anbehav.2016.03.025)

32. Schulman AH, Hale BE, Graves HB. 1970 Visual stimulus characteristics for initial approach response in chicks (*Gallus domesticus*). Animal Behaviour 18, 461–466.

33. Rugani R, Regolin L, Vallortigara G. 2010 Imprinted numbers: newborn chicks’ sensitivity to number vs. continuous extent of objects they have been reared with. Developmental Science 13, 790–797. (doi:10.1111/j.1467-7687.2009.00936.x)

34. Rosa Salva O, Mayer U, Vallortigara G. 2015 Roots of a social brain: Developmental models of emerging animacy-detection mechanisms. Neuroscience & Biobehavioral Reviews 50, 150–168. (doi:10.1016/j.neubiorev.2014.12.015)

35. Maekawa F, Komine O, Sato K, Kanamatsu T, Uchimura M, Tanaka K, Ohki-Hamazaki H. 2006 Imprinting modulates processing of visual information in the visual wulst of chicks. BMC Neuroscience 7, 1–13. (doi:10.1186/1471-2202-7-75)

36. Figler MH, Mills CJ, Petri HL. 1974 Effects of Imprinting Strength on Stimulus Generalization in Chicks (*Gallus gallus*). Behavioral Biology 545, 541–545.

37. Ham AD, Osorio D. 2007 Colour preferences and colour vision in poultry chicks. Proceedings of the Royal Society B: Biological Sciences 274, 1941–8. (doi:10.1098/rspb.2007.0538)

38. Jones CD, Osorio D, Baddeley RJ. 2001 Colour categorization by domestic chicks. Proceedings of the Royal Society B: Biological Sciences 268, 2077–2084. (doi:10.1098/rspb.2001.1734)

39. Osorio D, Vorobyev M, Jones CD. 1999 Colour vision of domestic chicks. 2959, 2951–2959.

40. Miura M, Nishi D, Matsushima T. 2019 Combined predisposed preferences for colour and biological motion make robust development of social attachment through imprinting. Animal Cognition (doi:10.1007/s10071-019-01327-5)

41. Wang M-S et al. 2020 863 genomes reveal the origin and domestication of chicken. Cell Res 30, 693–701. (doi:10.1038/s41422-020-0349-y)

42. Rosa-Salva O, Mayer U, Vallortigara G. 2019 Unlearned visual preferences for the head region in domestic chicks. PLoS ONE 14, 1–15. (doi:10.1371/journal.pone.0222079)

43. Zuk M, Ligon JD, Thornhill R. 1992 Effects of experimental manipulation of male secondary sex characters on female mate preference in red jungle fowl. Animal Behaviour 44, 999–1006. (doi:10.1016/S0003-3472(05)80312-4)

44. McGraw KJ et al. 2004 You Can’t Judge a Pigment by its Color: Carotenoid and Melanin Content of Yellow and Brown Feathers in Swallows, Bluebirds, Penguins, and Domestic Chickens. Condor 106, 390–395. (doi:10.1093/condor/106.2.390)

45. Mathis A, Mamidanna P, Cury KM, Abe T, Murthy VN, Mathis MW, Bethge M. 2018 DeepLabCut: markerless pose estimation of user-defined body parts with deep learning. Nat Neurosci 21, 1281–1289. (doi:10.1038/s41593-018-0209-y)

46. Versace E, Freeland L, Wang S, Emmerson MG. 2022 First-sight recognition of touched objects shows that chicks can solve the Molyneux’s problem. (doi:10.1101/2022.08.18.504388)

47. Schulman AH, Hale EB, Graves HB. 1970 Visual stimulus characteristics for initial approach response in chicks (Gallus domesticus). Animal Behaviour 18, 461–466. (doi:10.1016/0003-3472(70)90040-0)

48. Loconsole M, Pasculli MS, Regolin L. 2021 Space-luminance crossmodal correspondences in domestic chicks. Vision Research 188, 26–31. (doi:10.1016/j.visres.2021.07.001)

49. Kovach JK. 1971 Effectiveness of Different Colors in the Elicitation and Development of Approach Behavior in Chicks. Behaviour 38, 154–168.

50. Salzen EA, Lily RE, Mckeown JR. 1971 Colour prefrence and imprinting in domestic chicks. Animal Behaviour 19, 542–547.

51. Schaefer H, Hess H. 1959 Color Preferences in Imprinting Objects. Zeitschrift für Tierpsychologie 16, 161–172.

52. Gray PH. 1961 the Releasers of Imprinting: differential reactions to color as a function of maturation. Physiological Psychology 54, 597–601.

53. Bateson PPG, Jaeckel JB. 1976 Chicks’ preferences for familiar and novel conspicuous objects after different periods of exposure. Animal Behaviour 24, 386–39O. (doi:10.1016/S0003-3472(76)80048-6)

54. Hébert M, Versace E, Vallortigara G. 2019 Inexperienced preys know when to flee or to freeze in front of a threat. Proc Natl Acad Sci U S A 116, 22918–22920. (doi:10.1073/pnas.1915504116)

55. In press. A spontaneous gravity prior: Newborn chicks prefer stimuli that move against gravity | Zenodo. See https://zenodo.org/record/7437294#.Y-zaI8fP1D8 (accessed on 15 February 2023).

56. Lemaire BS, Vallortigara G. 2023 Life is in motion (through a chick’s eye). Anim Cogn 26, 129–140. (doi:10.1007/s10071-022-01703-8)

57. Johnson MH, Horn G. 1988 Development of filial preferences in dark-reared chicks. Animal Behaviour 36, 675–683. (doi:10.1016/S0003-3472(88)80150-7)

